# Simplest Model of Nervous System. II. Evolutionary Optimization

**DOI:** 10.1101/2023.11.24.568590

**Authors:** Anton V. Sinitskiy

## Abstract

In this work, we build upon a simple model of a primitive nervous system presented in a prior companion paper. Within this model, we formulate and solve an optimization problem, aiming to mirror the process of evolutionary optimization of the nervous system. The formally derived predictions include the emergence of sharp peaks of neural activity (‘spikes’), an increasing sensory sensitivity to external signals and a dramatic reduction in the cost of the functioning of the nervous system due to evolutionary optimization. Our work implies that we may be able to make general predictions about the behavior and characteristics of the nervous system irrespective of specific molecular mechanisms or evolutionary trajectories. It also underscores the potential utility of evolutionary optimization as a key principle in mathematical modeling of the nervous system and offers examples of analytical derivations possible in this field. Though grounded in a simple model, our findings offer a novel perspective, merging theoretical frameworks from nonequilibrium statistical physics with evolutionary principles. This perspective may guide more comprehensive inquiries into the intricate nature of neural networks.

## Introduction

How can we understand the nervous system regardless of specific molecular and cellular mechanisms and historical just-so-happened events? This challenge has persisted for a long time, largely because universal principles that could potentially apply to any nervous system (or their analogues in hypothetical extraterrestrial life forms) remain elusive. Likewise, we have yet to harness theoretical tools capable of distilling insights from these principles.

In this endeavor, we tackle a simple case of this quandary. Building upon the foundation laid in our preceding companion paper,^1^ we employ a simple model of the nervous system and its mathematical scaffolding. By formulating and subsequently resolving an optimization problem aligned with this model, we attempt to mirror the evolutionary optimization process. Despite the simplicity of the model, this approach offers certain predictions about the behavior and attributes of the modeled nervous system.

This investigation seeks to illuminate several overarching questions: Can we articulate broad-based predictions about the nervous system, uninfluenced by specific mechanisms or evolutionary history? To what extent can the principle of evolutionary optimization alone bolster our understanding and predictability of the nervous system’s attributes? What might mathematical derivations in this field look like?

Our exploration here is primarily confined to a simple model introduced in the previous paper.^1^ It is our aspiration that this endeavor, with its findings and solutions, will pave the way for broader and more intricate inquiries in this domain.^2^

## Methods

### Model definition

As in the previous companion paper,^1^ we consider a model of the simplest nervous system, inspired by a hypothetical scenario of the origin of the nervous system in Dickinsonia or similar animals from the Ediacaran period (henceforth referred to as “Dickinsonia” in quotes).^3^ Our model includes two dynamical variables: *x*, a generalized variable describing excitation of the nervous system, and *r*, the distance from the “Dickinsonia” to the nearest predator. The dependence of both variables on time *t* is described by the following stochastic differential equations:

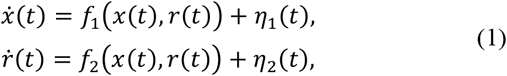

where 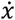 and 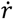 are the derivatives of *x* and *r* over time, *f*_1_ and *f*_2_ are some functions of *x* and *r*, and *η*_1_ and *η*_2_ are random variables (noise). Biologically meaningful expressions for *f*_1_ and *f*_2_ might be:

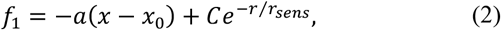

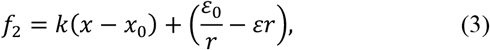

where *a, C, r*_sens_, *k, ε* and *ε*_0_ are positive constants, and *x*_0_ is a constant. The random variables η_1_ and η_2_ are assumed to obey Gaussian distributions with zero mean values and the following covariances:

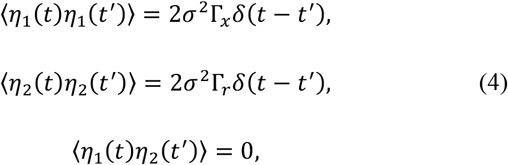

where Γ_*x*_ and Γ_*r*_ are positive constants, *δ*(·) is the Dirac delta function, and *σ*^2^ is a small parameter formally introduced to enable certain analytical derivations (note that it is not an independent parameter and could be absorbed by constants Γ_*x*_ and Γ_*r*_).

Recently, a powerful technique to analyze such dynamic systems has been proposed, inspired by methods from nonequilibrium statistical physics.^4-16^ Instead of considering individual stochastic trajectories that can be obtained from equations (1)-(4) by numerical integration, consider an ensemble of such systems characterized by a probability distribution function. In general, this function should be time-dependent; however, in many cases, a stationary probability distribution function *P*_*stat*_(*x,r*) exists. It is convenient to introduce a potential function *u*(*x,r*), an analogue of the common potential function in statistical physics generalized to nonequilibrium systems, defined from *P*_*stat*_(*x,r*) by the following equation:

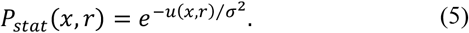

For the two-dimensional case under consideration, functions *f*_1_ and *f*_2_ from equation (1) can be expressed in terms of *u* as follows:^1^

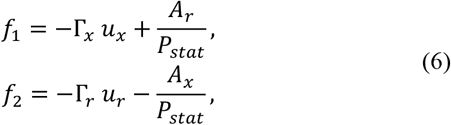

where *A* = *A*(*x,r*) is another function of *x* and *r*, and subscripts *x* or *r* here and below refer to partial derivatives with respect to the corresponding variable. In the literature, it is common to write these equations with the use of a function *Q* = *Q*(*x,r*) defined as:

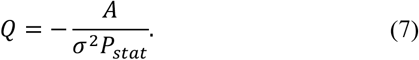

Then, equations (6) assume the following form:

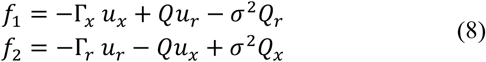

that effectively represents *f*_1_ and *f*_2_ in terms of the functions *u* and *Q*.

#### Novel approach: u and Q as independent variables

The power of the formalism presented above lies, among other things, in the fact that the *dynamics* of the system under study (the time derivatives of dynamic variables *x* and *r*, or more precisely, the deterministic components *f*_1_ and *f*_2_ of these derivatives) are expressed, by equations (6) or (8), in terms of the *time-independent* potential function *u*(*x,r*) and function *Q*(*x,r*) [or, alternatively, *A*(*x,r*)]. Furthermore, in the assumption of *ergodicity*,^17^ a time average along an infinitely long trajectory of an arbitrary function *F* that depends only on the current values of *x* and *r* can be computed as an average over the stationary distribution:

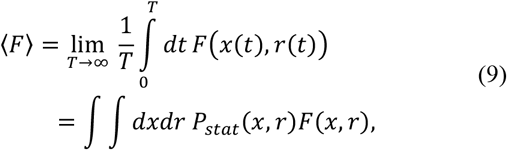

where the integration over *r* and *x* is performed over all their admissible values. The formalism outlined in the previous subsection is commonly used in the literature to analyze neural networks dynamics, given that functions *f*_*i*_ (and thus, *u* and *Q*) are predetermined.^4-16^

In this work, we use this formalism – for the first time, to the best of our knowledge – to address a new problem, namely, the problem of evolutionary optimization of neural networks. A traditional approach to this problem would be to define the specific forms for *f*_1_ and *f*_2_, for example, as in equations (2) and (3), integrate them to get an ensemble of trajectories, and then calculate the evolutionary fitness as defined in a given model as a temporal average over all trajectories. For dynamical equations of practical interest, integration is possible only numerically, and therefore, the optimization could scan a limited number of possible neural networks. By contrast, the approach presented in this work treats *u* and *Q* (or *A*) as independent entities, which define both the dynamics, namely *f*_1_ and *f*_2_, and the averages over the trajectories, with the use of equations (8) and (9), respectively. In this way, the expression for the evolutionary fitness can be obtained in an explicit, closed form, simplifying the task of its optimization. In next subsection, we address this problem of expressing the fitness for the introduced simple model of the nervous system in the form of a functional of *u* and *Q* (or *A*).

#### Cost of the nervous system functioning

The nervous system is an adaptation that requires the organism to expend additional resources, but ensures greater survivability. The evolutionarily optimal properties of the nervous systems are determined by the balance between evolutionary benefit and their cost. In generating an action potential, a neuron loses some of its ions (for example, K^+^ ions), while additional amounts of other ions (for example, Na^+^ or Cl^−^ ions) penetrate the cell. To maintain the neuron’s ability to generate action potentials, ions need to be moved back, against their electrochemical potentials, a job performed by ion pumps consuming ATP. There is a simple proportion between the number of pumped ions and the number of ATP molecules spent, which allows estimating the cost of the nervous system’s work by the number of ions moved through neuron membranes by ion pumps.^18-23^ The number of ions to be pumped equals the number of ions *q* passed through the neuron membrane during the generation of action potentials, which, in its turn, can be found as the integral over time of the current *i*(*t*) through the membrane. Using the standard assumption that the membrane resistance, *R*_*mem*_, is constant, the current can be expressed through the difference of the membrane potential *x* from the resting potential *x*_0_. This leads to the following expression for the number of ions passing through the membrane during the time *T*:

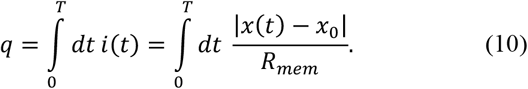

A typical situation is the depolarization of a neuron, where *x* > *x*_0_. However, hyperpolarization is also possible (for example, when excessive amounts of Cl^−^ ions enter the neuron in inhibitory synapses). In this case, *x* < *x*_0_, but returning the neuron to the resting state also requires the work of ion pumps, still spending positive amounts of free energy, which explains why we included the absolute value of *x* – *x*_0_ on the right-hand side of equation (10). Thus, over a sufficiently long period of time *T*, the work of the neuron requires the expenditure of a number of ATP molecules proportional to *q*, and therefore, proportional to 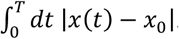 Unlike ion flows through the membrane, the work of ion pumps is roughly even over time. Thus, the average cost of the ion pumps’ work per unit time, over a sufficiently long period *T*, is proportional to the temporal average of |*x*(*t*) − *x*_0_|.

Note that more general thermodynamic considerations^24-26^ can be used to derive the cost of the nervous system functioning, without references to specific details like the stoichiometric calculation of the number of ATP molecules. In particular, in our model, such a derivation leads to the same final result. The work performed by the system to move an infinitely small charge *dq* against the electrochemical gradient equals (*V-V*_*eq*_)*dq*, where *V* is the current otential and *V*_*eq*_ is the equilibrium potential value at given concentrations. Since the number of ions passing through the membrane during one action potential barely changes the ion concentrations inside and outside the cell, and since ion pumps operate at the resting potential most of the time, the value of (*V-V*_*eq*_) is constant, which means that the work performed by the system (the cost of a neuron’s functioning) is proportional to 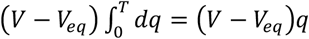, hence, on average proportional to |*x*(*t*) − *x*_0_|, the same result as above.

Assuming ergodicity of the system, we conclude from equation (9) that the cost of the nervous system’s functioning (in units of free energy per unit time) is proportional to the value of *I*_1_ defined as

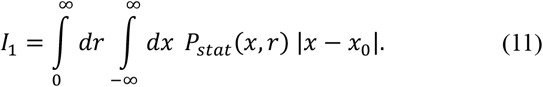

Note that this cost includes the work cost of sensors. The mechanism of sensor action typically involves ion channels that open in response to external signals, leading to changes in the membrane potential of the sensor neuron changes due to ion passage. As with the generation of a spike, the expenditure of free energy is associated with the reverse pumping of ions against the gradient of the potential, and therefore, was taken into account in the calculation above.

#### Cost of the work of effectors

The cost of the nervous system computed in the previous subsection does not include the work cost of effectors (for example, cilia, whose coordinated beating presumably allowed Dickinsonia to move in space). As known from classical mechanics, the work performed during mechanical movement equals the product of force *F* and speed *v*. Further, when moving in a viscous medium, the movement speed is directly proportional to the force, *v* ∼ *F*, which means that the work per unit of time is proportional to *v*^2^. The movement speed of “Dickinsonia” *v* in this model can be found as the difference Δ*f*_2_ between the speed at which the distance from “Dickinsonia” to the predator *f*_2_ changes, and the contribution to this speed from the motion of the predator itself *f*_*pred*_(*r*) = *ε*_0_*/r* – *εr* that was calculated earlier:^1^

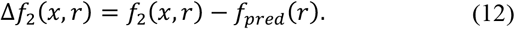

Here we assume that the predator is moving towards “Dickinsonia”, and “Dickinsonia” is running away from it, i.e., the speeds are directed in opposite directions. For example, with the expression for *f*_2_ given by equation (3), Δ*f*_2_(*x, r*) = *k*(*x* − *x*_0_), and therefore, the cost of the work of effectors is proportional to the average of *k*^2^(*x* − *x*_0_)^2^.

As in calculating the cost of a neuron’s functioning, we replace the time average by the ensemble average, and conclude that the average cost of actors engaged by the nervous system is proportional to the average value of 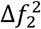, that is, to *I*_2_ defined as

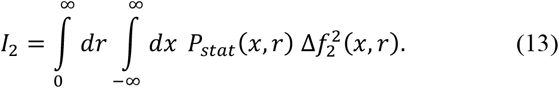

#### Probability of an organism’s death

As demonstrated in Appendix A, in the stationary state, the probability *P*_*death*_ of “Dickinsonia” being “eaten” (critically damaged by external digestion by another “Dickinsonia”) per unit of time can be found as

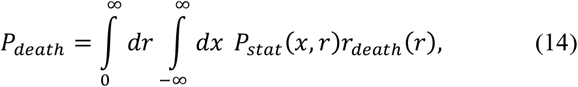

where *r*_death_(*r*) is the probability of a “Dickinsonia” being eaten if the nearest predator is at a distance *r*. In the realistic case of a nervous system – not necessarily perfectly optimized, but nevertheless working more or less efficiently to evade predators – the stationary distribution *P*_*stat*_ will have only a light tail in the region of the values of *r* where predation is possible. This justifies the use of the distribution *P*_*stat*_ derived in the context of infinitely long trajectories to estimate *P*_*death*_ defined in the context of finite, but sufficiently long [in comparison to the timescales set by equations (1)] trajectories.

#### Evolutionary optimality in general

The population growth rate of “Dickinsonia” *R*_0_ in the absence of predators – and hence the absence of the need to expend resources on evading them – is determined by the availability of bacterial mats and other factors not related to the predator evasion by the nervous system. Within the framework of this model, we will consider this value *R*_0_ as constant. The appearance of predators leads to the death of some of the “Dickinsonia”, thus reducing the growth rate of the population by the amount of *P*_*death*_, which can be found from equation (14). Furthermore, the functioning of the nervous system and effectors, while reducing *P*_*death*_, also requires the expenditure of “Dickinsonia” resources, which can no longer be directed towards population growth, thus reducing the growth rate by amounts proportional to *I*_1_ and *I*_2_. Thus, the population growth rate *R* in the presence of predators and with the functioning nervous system can be calculated as:

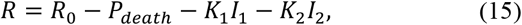

where *K*_1_ and *K*_2_ are positive constants. Natural selection optimizes functions *f*_1_ and *f*_2_, among those that are physically and biologically achievable, to maximize the value of *R*, or equivalently, minimize the functional *I* defined as

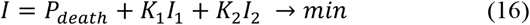

Note that the values of *P*_*death*_, *I*_1_ and *I*_2_ are non-negative, therefore:

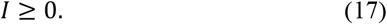

The functions *f*_1_ and *f*_2_ are uniquely defined by functions *u* and *Q* by equations (8), and therefore, the functional *I* can be considered as a functional solely of *u* and *Q*:

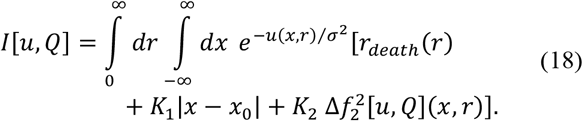

This representation is more convenient since expressions for *P*_*death*_, *I*_1_ and *I*_2_ were obtained above in terms of *P*_*stat*_ (that is, *u*) and *Q*, not *f*_1_ and *f*_2_. The notation Δ*f*_2_[*u,Q*] means the same as just Δ*f*_2_; by adding [*u,Q*] to the notation, we emphasize that function *f*_2_, and hence Δ*f*_2_, is set by the choice of *u* and *Q* according to equation (8).

When minimizing *I*, in general, certain restrictions must be imposed on permissible values of *u* and *Q*, reflecting physically and biologically possible functions *f*_1_ and *f*_2_. In particular, the structure of the external physical world cannot be changed by natural selection acting on “Dickinsonia” in this model (we neglect large-scale ecological processes). Therefore, the deterministic component of *f*_2_ with the nervous system not working (*x* = *x*_0_) should be determined by external physical factors, and

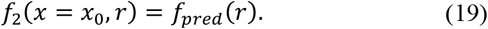

Also, *P*_*stat*_ must be normalized to unity, implying:

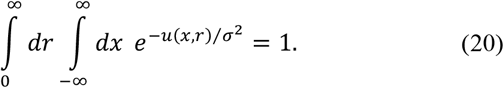

As discussed below, some additional constraints on *u* and *Q* may exist (see Results).

## Results

### Set of principles is incomplete for an ab initio derivation of the optimal solution

Naively, the functional *I* given by equation (18), under constraints (19) and (20), could be minimized to zero [which, according to equation (17), is its minimal value in principle] in the following way. Note that *I*[*u,Q*] depends on *Q* only via the third term in the integrand, and this term is non-negative, because *K*_2_ > 0 and 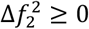. Therefore, the contribution to *I* from this third term can be minimized – namely, made equal to zero – if *Q* is chosen in such a way that Δ*f*_2_(*x,r*) = 0 at all values of *x* and *r*. The first two contributions to *I* in this case are not affected, assuming that *u* does not change, because, as follows from equation (18), these two terms do not depend on *Q*. The value of *Q* that satisfies this condition can be found from equations (6) and (7), resulting in:

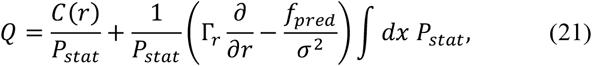

where *C*(*r*) is an arbitrary function. Note that restriction (19) in this case is automatically satisfied by construction. Furthermore, the first two contributions to the functional *I* can be put to zero by choosing the potential *u* in the following form:

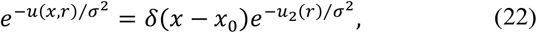

where *δ*(·) is the Dirac delta function, and *u*_2_ is any function of *r* that goes to +∞ at all *r* for which *r*_death_(*r*) > 0. As a result, *I* = 0.

However, this solution is biologically unmeaningful. In particular, nullification of Δ*f*_2_(*x,r*) not only at *x* = *x*_0_, as demanded by condition (19), but also for all other *x* implies that the effector is not controlled by the nervous system, and the temporal dynamics of *r* depends only on the environment (movement of predators), but not on the organism under consideration, which, evidently, cannot be the global evolutionarily optimal solution for a nervous system.

It is not entirely clear to us yet what kind of additional restrictions could prevent solution (21). We hypothesize that it may be related to general conditions on the existence of the stationary distribution *P*_*stat*_, which is required for the validity of all formulas in this paper, starting from equation (5). Though the choice of *f*_2_ as *f*_2_(*x,r*) = *f*_*pred*_(*r*) should not lead to divergences of the trajectories in time, this may be not true for *f*_1_ corresponding to *Q* and *u* set by equations (21) and (22). We leave this question for further work.

As of now, we can only state that the set of principles given by equations (18)-(20) is incomplete for an *ab initio* derivation of the evolutionarily optimized properties of a nervous system, because it does not exclude biologically nonsense solutions like those given by equations (21) and (22).

### Analytical results on evolutionary optimization

To circumvent this incompleteness problem, we now restrict our consideration to such *u* and *Q* that correspond to biologically and physically plausible dynamics of the system. Based on our results from the previous paper,^1^ we assume that the potential function has the following form:

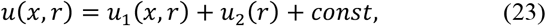

where

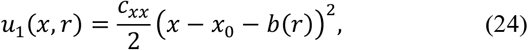

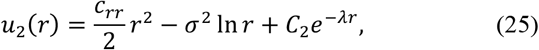

where we used a shorthand notation

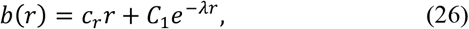

and *c*_*xx*_, *c*_*rr*_, *c*_*r*_, *C*_1_, *C*_2_ and *λ* are positive constants. Note that, in terms of the notation from the previous paper, we chose *c*_*ln*_ = 1 and the same value *λ* for the constants in the exponential functions in equations (25) and (26) to make derivations simpler.

As shown in Appendix B, with the proper choice of the function *Q*, which is possible with any values of the above-listed parameters in *u*(*x,r*), the speed of escaping the predator is directly proportional to the activity of the nervous system:

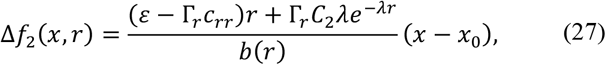

and, in particular, in the absence of such activity condition (19) is satisfied. Thus, as soon as the fraction on the right-hand side of equation (27) does not vanish, the problem of the loss of connection between the nervous system and the effectors, as in the previous subsection, does not emerge.

Using this approach, the general functional *I*[*u,Q*], which depends on functions *u* and *Q*, is simplified to depend only on the numerical parameters *c*_*xx*_, *c*_*rr*_, *c*_*r*_, *C*_1_, *C*_2_ and *λ*:

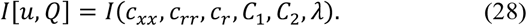

Thus, varying the values of *c*_*xx*_, *c*_*rr*_, *c*_*r*_, *C*_1_, *C*_2_ and *λ* (all of which are assumed positive) provides a particular case of evolutionary optimization of the primitive nervous system of the “Dickinsonia”, ensuring that the resulting dynamics is biologically meaningful. Note that Γ_*x*_ and, to a lesser extent, Γ_*r*_ may also be adjusted by the evolution, by means of changing the level of the noise in the neuron and the effector, though the contribution to Γ_*r*_ from the noise in the dynamics of the environment is not affected. However, in this work, we consider Γ_*x*_ and Γ_*r*_ to be constants.

Plugging specific expressions (23) and (27) into equation (18), we note that integration over *x* in equation (18) can be performed analytically (Appendix B), leading to:

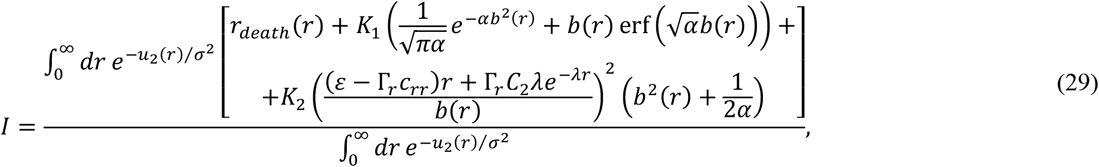

where *α* = *c*_*xx*_ ⁄2*σ*^2^.

The key result of this section is that the value of *I* can be made arbitrarily small by the proper choice of the parameters *c*_*xx*_, *c*_*rr*_, *c*_*r*_, *C*_1_, *C*_2_ and *λ*. Specifically, in the limit of *c*_*xx*_ → +∞ (and hence *α* → +∞), the right-hand side of equation (29) tends to

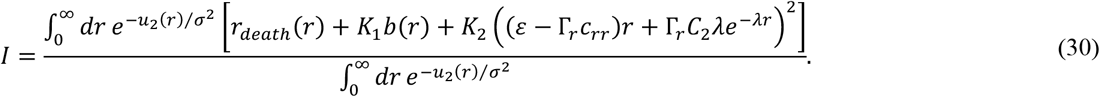

Then, in the limits of *c*_*r*_ → 0^+^ and *c*_*rr*_ → *ε*/Γ_*r*_, the functional *I* further simplifies to:

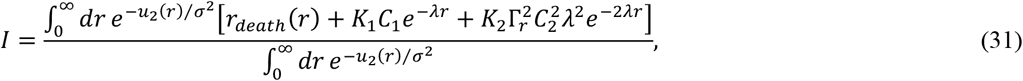

or, in an expanded form,

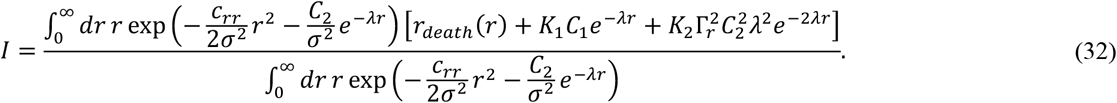

Finally, by increasing *C*_2_ to +∞ while maintaining *λ* as a finite positive constant, the region with large distribution values shifts towards larger *r* values. This results in decreased values of *σ*^−*σr*^ in that region, reducing the part where *r*_*death*_(*r*) is non-zero. As a result, *I* → 0 is attainable in the combination of the limits of *c*_*xx*_ → +∞, *c*_*r*_ *σ* 0^+^, *c*_*rr*_ *σ ε*/Γ_*r*_, *C*_2_ → +∞ and *λ σ const* > 0 (as for *C*_1_, it may stay constant or grow to infinity as *C*_2_).

What dynamic equations of the form given by equations (1)-(3) would correspond to these limits? As demonstrated in the previous paper, the potential given by equations (23)-(25) is, strictly speaking, only an approximation to the exact potential corresponding to dynamical equations (1)-(3), so we can expect to get a qualitative guess, not an exact prediction. Nevertheless, we try to make such a prediction, with a consequent check by numerical integration of the dynamical equations in next subsection. As we can see from the relationships between the coefficients in the potential *u*, equations (23)-(26), and the coefficients in dynamical equations (1)-(4), derived in the previous paper and duplicated below:

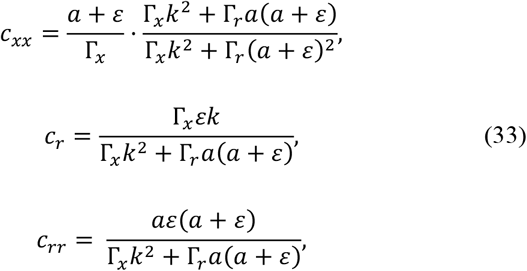

given the constant values of *ε*, Γ_*x*_ and Γ_*r*_, the limit of *c*_*xx*_ → +∞ can be achieved only if *a* → +∞, while *k* might stay finite or go to infinity slower or faster than *a*. In all these cases, *c*_*r*_ → 0^+^, as desired. However, the limit of *c*_*rr*_ → *ε*/Γ_*r*_ can be achieved only if *k* stays finite or goes to infinity slower than *a*. Finally, given the qualitative connection between the terms *C*_1_*e*^−*λr*^ and *C*_2_*e*^−*λr*^ in the approximate potential and the sensory 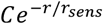 term in the dynamical equations, we expect that the limit of *C*_2_ → +∞ corresponds to *C* → +∞ in the sensory term in equations (1)-(4). As a result, we arrive at the dynamical equations that can conventionally be written as:

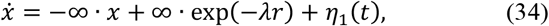

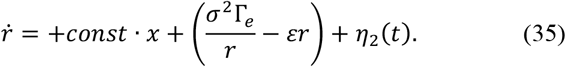

Evidently, the infinite values of the two prefactors in a biological system are unachievable. In practical terms, the numerical values of these prefactors will be constrained by the finite timescales associated with molecular mechanisms, such as ion channel functional states transitions and intercellular signal transduction. Also, after a sufficiently advanced minimization of the functional *I*, other components of the cost may become important, preventing it from approaching the zero value. Nevertheless, this theoretical result is valuable from the viewpoint of trend prediction. It demonstrates that evolutionary optimization may lead to a low working cost as long as the nervous system is already in place (note that *I* does not include the fixed costs of a nervous system, such as structural, genetic and metabolic costs in the absence of active firing). This result may guide our understanding of ion channels and sensory mechanisms in the way that the global optimization of the evolutionary fitness manifests in local evolutionary optimization for properties like sensitivity and kinetics of ion channels. Also, this result is important from the conceptual viewpoint, because it supplements a common perception of a nervous system as evolutionarily expensive^18,20,21,27-29^ with an understanding that this cost may be pretty efficiently optimized in comparison to preadaptive states.

Finally, note that the derivation presented in this subsection seems robust, and we expect that it may stay valid for a wider range of functional forms for *r*_*death*_(*r*) and expressions for the cost components *I*_1_ and *I*_2_. In order for the contribution from *r*_*death*_(*r*) to *I* to vanish in the limit of *C*_2_ *→* +∞, *r*_*death*_(*r*) needs only to be localized in the region of small *r* (or, in a more general case, the region of values of *r* that the organism evades in the corresponding limit), and it is not required for it to following specific conditions given as examples in Appendix A, like being stepwise, constant at *r* < *r*_*death*_, etc. Moreover, minimizing the cost component *I*_1_, by narrowing the distribution over *x* as achieved in the *c*_*xx*_ *→* +∞ limit, is not limited to the expression in equation (11); it can also apply to its generalizations with a non-linear dependence on *x*. The only necessary condition is that the integrand equals zero at *x* = *x*_0_, that is, when the neuronal membrane is at its resting value, which seems to be quite general. Finally, similar considerations may be extended to *I*_2_. We expect that a proper choice of limits for the parameters in *u*(*x,r*) may nullify *I*_2_ even for more complex, non-linear dependencies of Δ*f*_2_(*x,r*) on *x*, taking into account that they should vanish in the limit of small *x* – *x*_0_ (whether linearly, as in the analyzed case, or non-linearly) due to condition (19). Hence, we expect that the conclusion on the evolutionary optimization of a primitive nervous system by shortening the timescale of its activation and increasing its sensitivity may stay valid beyond the specific model introduced above.

### Numerical results on optimized functions

To illustrate these theoretical derivations and check the validity of the used approximations [ergodicity, approximate *u* and matching of approximate results to dynamical equations (34)-(35)], we ran numerical simulations of equations (1)-(4) with parameters *a* and *C* scaled from their values in the previous companion paper^1^ by the same factor *s*:

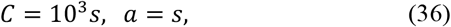

while the other parameters (including *k*) were kept unchanged. The values of *s* were increased as powers of two. At each value of *s*, the generated trajectories [*x*(*t*),*r*(*t*)] were used to compute the values of *P*_*death*_, *I*_1_ and *I*_2_ as temporal averages of *χ*(*r*), |*x* – *x*_0_| and *k*^2^(*x* – *x*_0_)^2^, respectively; for *χ*(*r*), the critical distance of *r*_*death*_ = 5 was chosen to ensure plausible (small, but not negligible) values of *P*_*death*_. The length of the simulations was chosen in such a way that the resulting values of *P*_*death*_, *I*_1_ and *I*_2_ were numerically stable, which limited the maximal value of *s* to 64 (greater *s* require smaller integration steps and at the same time longer trajectories for accurate estimates of the computed values, especially *P*_*death*_).

As these simulations demonstrate, each of the three components of the evolutionary cost of the model nervous system decreases with increasing the scale *s* (Fig. 1). The fraction of the organisms in the dangerous zone drops from 0.15% for the initial values of the parameters to 0.04% after increasing *a* and *C* just by a factor of *s* = 2, and continues rapidly dropping with the subsequent increase in *s* (Fig. 1, left scale). Both the integrals *I*_1_ and *I*_2_ also decrease with the increasing *s* (Fig. 1, right scale), though this decrease is more gradual than in the case of *P*_*death*_. Hence, the total evolutionary cost of the working nervous system in this model *I* consistently decreases with rescaling *a* and *C*, and this conclusion holds true regardless of specific values of the conversion coefficients *K*_1_ and *K*_2_ in equation (16).

**Fig. 1.**
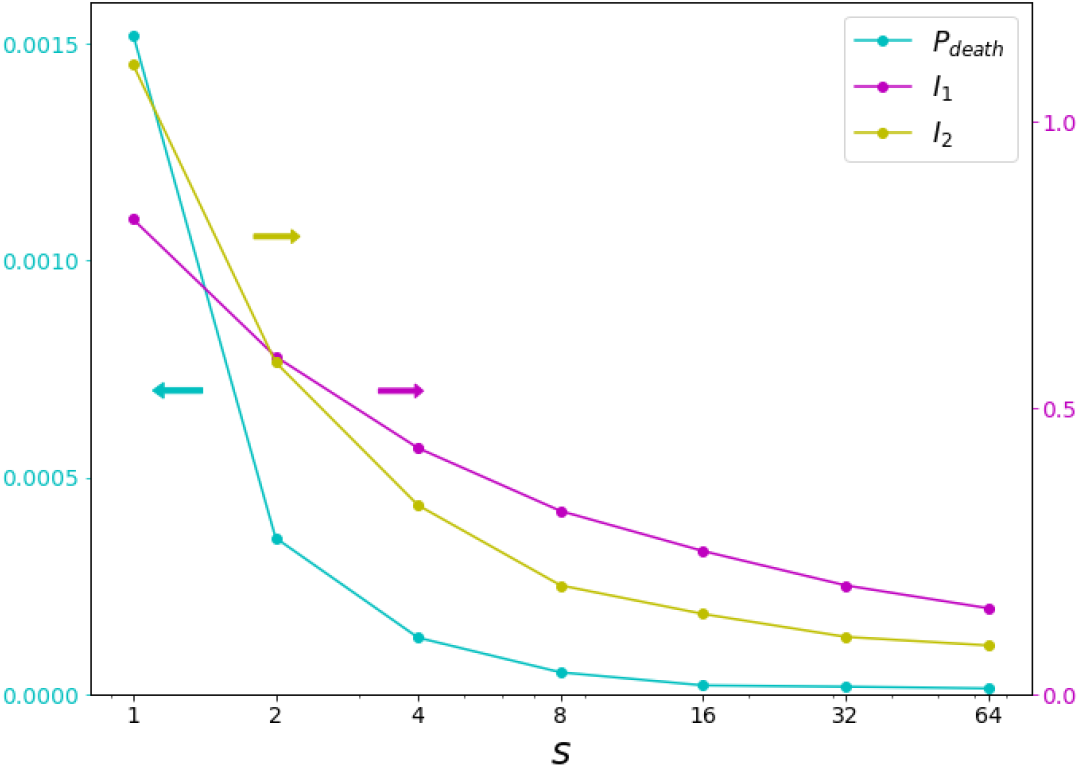
Rescaling the parameters *a* and *C*, as theoretically predicted, leads to a decreasing evolutionary cost of the nervous system. The scaling parameter *s* equal to 1 corresponds to the original values of the parameters *a* and *C* in the previous work.^1^ Larger values of *s* correspond to the nervous system more sensitive to the sensory signal and faster returning to the resting state. One of the components of the evolutionary cost, *P*_*death*_, refers to the left vertical scale (*cyan*), while the other two components refer to the right vertical scale (*magenta, I*_1_, *yellow, I*_2_).

Visual investigation of a typical trajectory generated with parameters *a* and *C* rescaled with *s* = 64 reveals significant changes in comparison to the original preadaptive state reported in the previous accompanying paper^1^ (Fig. 2). The optimized model generates more pronounced spikes of neural activity, with much higher signal-to-noise ratio and narrower peaks. As stated earlier,^1^ the original model, given its simplicity, could be interpreted as corresponding to a preadaptive state of an early nervous system, with a noisy unreliable functioning, evolutionarily advantageous only on average, over a large number of events (Fig. 2a,b). The change of the neural activity pattern after the parameter rescaling (biological interpretation: after evolutionary optimization) matches the expectations of a more reliably functioning nervous system and a better detection of each event of a predator approach (Fig. 2d,e). While before optimization, multiple false positive and false negative events of neural excitation were observed (the nervous system excites in the absence of a predator nearby or, vice versa, does not excite when a predator approaches, Fig. 2a,b), the optimized model nervous system does not allow for such false positive or negative events. On longer timescales, the optimized model demonstrates a greater preference for depolarization events over hyperpolarization events (Fig. 2f) in comparison to the original model (Fig. 2c), reflecting a stronger role of depolarization events in the detection of evolutionarily important signals from the environment. Thus, the proposed model, despite its simplicity, captures the most important aspects of the evolutionary optimization of a primitive nervous system, in particular, the emergence of its spiky dynamics.

**Fig. 2.**
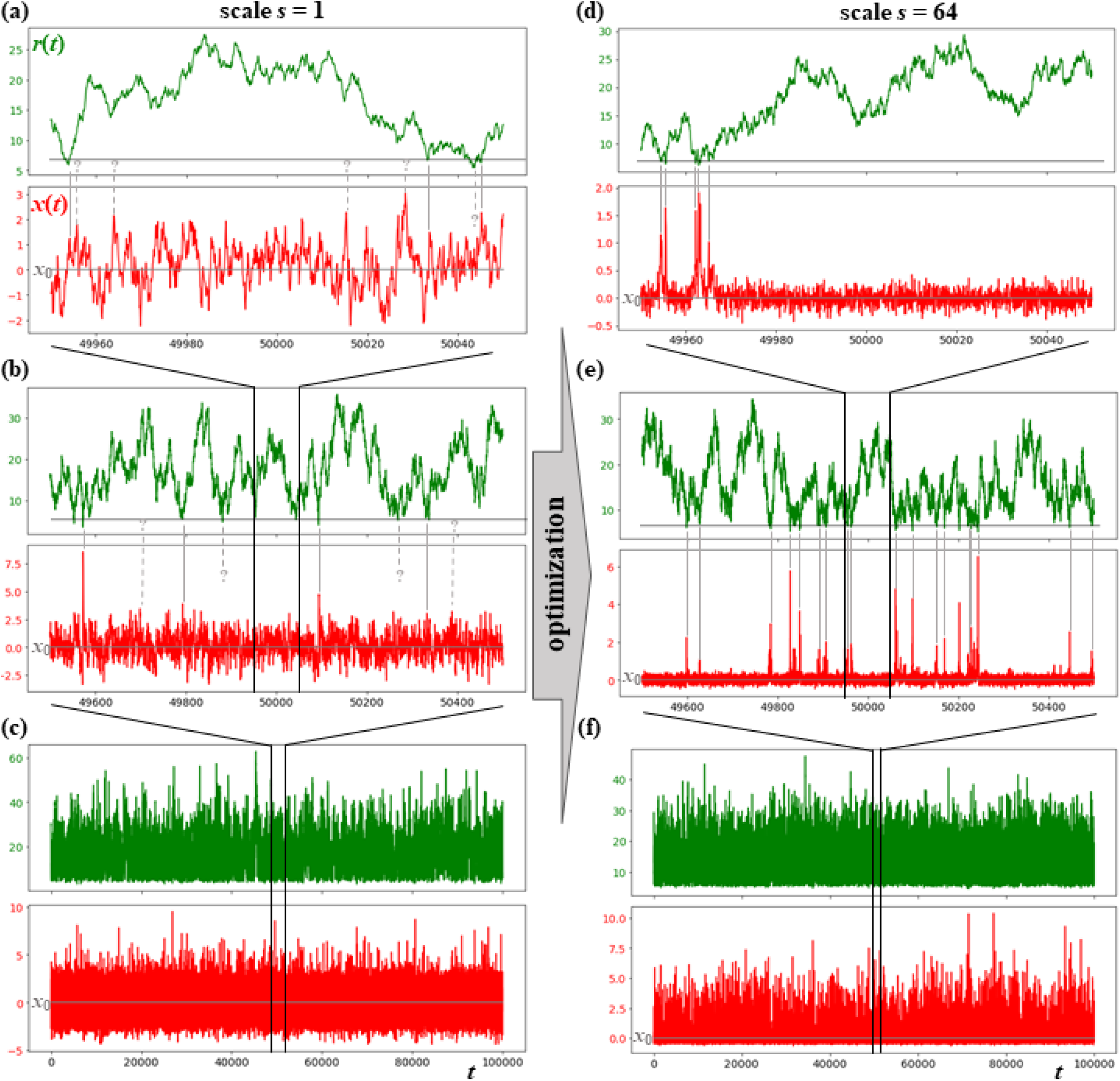
Rescaling the parameters *a* and *C* leads to a more evolutionary optimized dynamics of the model nervous system (*right panels*), as compared to the preadaptive state (*left panels*). On shorter timescales, the optimized model demonstrates much lower signal-to-noise ratio (d,e) for the nervous system activity *x* (*green*) than the original model (a,b), while the noise in the environmental variable *r* (*red*) remains comparable. After optimization, the events of neuronal excitation strictly coincide with the events of predator approach (d, e, *grey vertical lines*), while in the pre-optimized state, multiple cases of predator approach without neuronal excitation or vice versa were observed (a, b, *dotted vertical lines with question marks*). Also, optimization results in narrower peaks of neural activity (d), which can be conditionally interpreted, given the simplicity of the model, as a more ‘spiky’ behavior (as opposed to a graded potential response) than in the original model (a). On longer timescales, stationary distributions over *x* and *r* are reached in both cases (c,f). Note that depolarization is more likely to occur than hyperpolarization in both cases, but the asymmetry is much more pronounced in the optimized model (f).

## Discussion

In this work, we present a specific example of how the formalism from nonequilibrium statistical physics could be coupled with an evolutionary approach to make general predictions about the nervous system. This union enables us to express both dynamics and statistics of the studied system (a simple nervous system in a simple environment) using a common set of entities, namely functions *u* and *Q* as defined in Methods. Building on the idea that existing organisms operate close to their evolutionary optimum, we demonstrated how predictions about the nervous system can be done in this framework without resorting to specific molecular and cellular mechanisms or events in the evolutionary history.

The central result of this work is a general derivation of certain characteristics of the “Dickinsonia” nervous system, specifically, the pronounced neuron excitation and relaxation, and the high sensitivity of the nervous system to an approaching predator. Our model suggests that these behaviors are direct consequences of broad evolutionary factors and optimization of nervous system function. While these results are rooted in mathematical derivations, we also reinforced their validity through numerical simulations.

While the presented model, based on a hypothetical scenario of the emergence of a primordial nervous system, has its particularities, its underlying formulations, especially the expressions for fitness components (*P*_*stat*_, *I*_1_ and *I*_2_), seem fairly general. We expect that even if the conditions on these functionals were loosened, evolutionary optimization would lead us to qualitatively similar results.^2^

Even though we previously warned against naïve intuitive conclusions,^1^ here we will try to present an intuitive interpretation of the general solution derived in this work. Hopefully, it will help us develop the right intuition for this type of problems for the future. Starting from uncoupled variables *x* and *r* (nervous system that does not feel the external world and the external world that is not changed by the nervous system), we aim to infer the evolutionarily optimal solution to the problem. In this initial uncoupled state, *x*(*t*) follows the equation 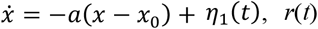 follows the equation 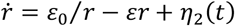, and therefore, their stationary distribution probability density will factorize: 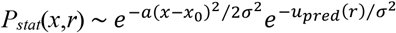, where *u*_*pred*_ was derived in Appendix A of the previous paper^1^ as the effective potential for the nearest predator random distribution. In the limit of *a →* +∞, *I*_1_ and *I*_2_ vanish, but *P*_*death*_ stays relatively high (no predator evasion leads to a high probability of being eaten). To minimize *P*_*death*_, it is necessary for the probability density *P*_*stat*_(*x,r*) at *r* < *r*_*death*_ to become small by redistributing a part of the probability to the region with *r* > *r*_*death*_. But in this case, near the border of *r* = *r*_*death*_, *P*_*stat*_ increases more than for a random predator distribution, therefore, *u* decreases more, and thus there is a large negative additional contribution to partial derivative *u*_*r*_ relative to the partial derivative *u*_*pred,r*_ of the undisturbed effective potential *u*_*pred*_. As a result, taking into consideration the relationship between *f*_2_ and *u*_*r*_ given by equation (8), condition (19) is no longer met and a significant contribution to *I*_2_ may appear due to positive values of 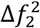. How can this contribution be avoided? In equation (8), the expression for *f*_2_ also includes terms with *Q*, and thus the change in *u*_*r*_ can be compensated by appearance of a non-zero value of *Q*, which is quite logical, since the operation of the nervous system (unlike the initial uncoupled state) is a dissipative process. Note that in the limit of *a →* +∞, infinitely small *Q* should have a finite effect, since the partial derivative *u*_*x*_ should be infinitely large for infinitely narrow distributions over *x*, which is required to minimize *I*_1_. The terms with partial derivatives of *Q* in equation (8) in this limit can be neglected, because they are multiplied by a finite value of *σ*^2^. Remarkably, together with the additional contribution to *f*_2_, this non-zero value of *Q* also, according to equation (8), results in a new contribution to *f*_1_, which leads to a dependence of the dynamics of the neuronal membrane potential also on the distance to the predator *r*, that is, to the emergence of the sensory role of the neuron, completing the picture of the optimal solution.

As with all models, our model comes with its limitations. It is built upon the conceptual simplifications elucidated in our preceding companion paper.^1^ Most importantly, we emphasize a holistic understanding of the nervous system. At the same time, to simplify it to an acceptable degree, we kept the model’s dimensionality minimal – just one variable for the nervous system and one for the state of the environment. Additionally, we made the model as independent as possible from specific molecular or cellular mechanisms. These simplifications allow us to emphasize the core components of a model of a nervous system: treatment of the nervous system in the context of its environment and the pivotal role of evolution in dictating its properties. These simplifications pave the way for a more accessible, albeit stripped-down, platform to conduct studies of general problems about the nervous system in a more tractable (including comprehensive numerical simulations) case.

From a technical perspective, our model incorporates some additional limitations. The zero limit values for both the probability of being preyed upon and the energy cost of neurons and actors cannot be taken literally: in the real world, a specific material implementation will not allow the nervous system to reach this limit. The assumption of ergodicity may be broken if the potential *u* presumes disconnected parts of the phase space^17^ (for example, note that the long-distance limit solution for *u* given in the previous paper allows for negative values of *r*, while the corrected solution contains a prohibitively high barrier as *r →* 0^+^; however, a formal mathematical solution could contain a potentially allowed region with negative *r* that trajectories starting from positive *r* will not achieve on reasonable timescales). Also, while the model predicts sharp peaks in neuronal activity and suggests that evolutionary optimization should make them even sharper, these peaks, strictly speaking, cannot be interpreted as spikes. Known models of spiking neurons either require more than one dimension (e.g., the Hodgkin–Huxley^30^ or FitzHugh– Nagumo^31,32^ models) or include aspects incompatible with the Fokker-Planck formalism we employ (e.g., a discontinuous reset of the potential in various integrate-and-fire models,^33,34^ or a periodic nature of the theta variable in the theta model^35^). Therefore, the difference between spiking and non-spiking (graded potential) neurons is not thoroughly captured by our model, and a discussion has to restrict to the difference of sharp and narrow vs. diffuse and wide peaks of neuronal activity.

Our model opens numerous avenues for exploration. Future studies could consider additional constraints on the functions *u* and *Q*, accounting for physical and biological realities and solving the problem of the incomplete set of restrictions, as described in Results. In particular, the intuitively plausible expressions for *f*_1_ and *f*_2_ given in equations (2) and (3), assume a separation of contributions from variables *x* and *r*, which, however, is not automatically ensured in the general case given by equations (6) and (8). Due to the presence of terms involving *Q* in these equations, separation of variables in *f*_1_ and *f*_2_ does not imply their separation in *u*(*x,r*). We began investigating the relationship between such separability and the assumption of constant *Q*, but this work is still ongoing. Another possible direction would be to revise the expressions for the cost of the nervous system operation, in particular, identifying the terms that may become prevailing during the minimization of *I*_1_ and *I*_2_ and their approach to zero. On the one hand, such attempts may be biology-guided, based on more detailed analysis of specific molecular mechanisms in existing organisms. On the other hand, these efforts may proceed in the direction of a more abstract treatment, establishing a connection to more general physical concepts and expressions for the ‘cost’ of a nonequilibrium dissipative system (e.g., entropy production rate).^36^

In conclusion, in this work we propose a simple (perhaps the simplest) model of the nervous system that may serve as an initial step, offering a novel lens to perceive general characteristics of the nervous system and bridging nonequilibrium statistical physics with evolutionary principles. The complexities and nuances of the nervous system demand an integrative approach, and we hope that our study propels us closer to unraveling them.

## Appendices

### Appendix A. Probability of death expressed in terms of the stationary probability distribution

The time-dependent probability distribution function *P*(*x,r,t*) for the ensemble of systems described by dynamic equations (1) and (4) obeys the Fokker-Planck equation:^1^

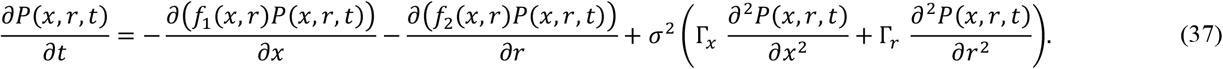

Let us rewrite this equation in terms of the total number of organisms *N*(*x,r,t*) in state *x,r* at time *t*. Initially, we assume that the total number of organisms at time *t* is constant; therefore *N*(*x,r,t*) is proportional to the probability density *P*(*x,r,t*), hence

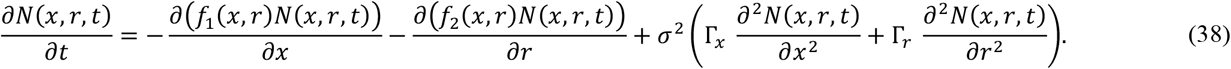

The obtained equation describes the change in the number of organisms in each of the states *x,r* due to the temporal dynamics of the states of each of the organisms according to equation (1). Now let us set aside the assumption that the total number of organisms remains constant and add to the right side of the equation another term that accounts for the death of organisms:

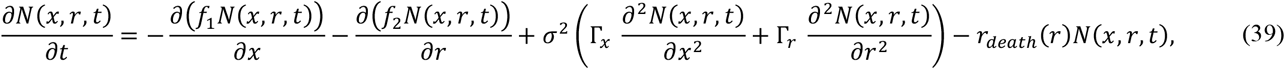

where *r*_death_(*r*) is the probability of a “Dickinsonia” being eaten by a predator located at a distance *r*. In this simplest model, we assume that this probability does not directly depend on the state of the victim’s nervous system *x*, but only on its position in space relative to the predator. In this case, the total number of a “Dickinsonia” at time *t* can be found by integrating over all possible states *x,r*:

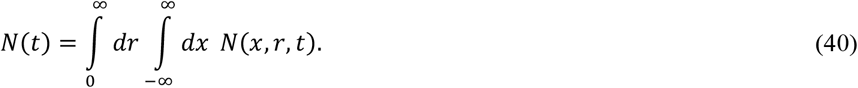

The rate of change of the total number *N*(*t*) over time can be calculated by integrating equation (39):

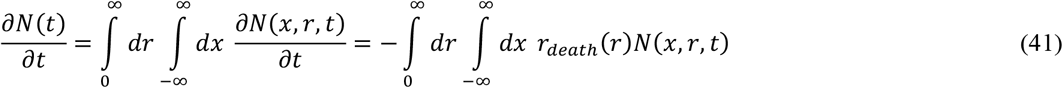

(the contributions of the terms that were present in the original Fokker-Planck equation cancel out, because the original equation (38) preserved the total number of organisms). Returning from the description in terms of the number of organisms to the description in terms of probability distributions, we find that the total probability *P*_*death*_(*t*) of a “Dickinsonia” being eaten per unit of time can be expressed as follows:

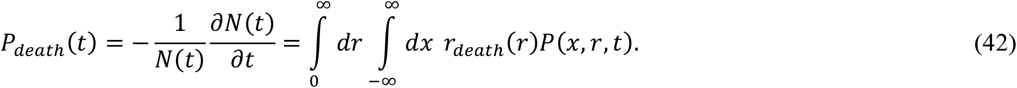

Then, in the stationary state, this probability *P*_*death*_ of being eaten per unit of time becomes time-independent and can be calculated as

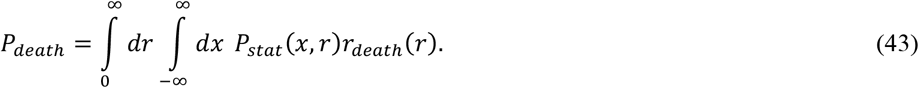

If we assume that the probability of being eaten by the predator diminishes beyond a certain threshold distance *r*_*death*_ and remains constant below that distance, then

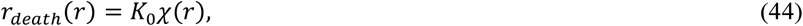

where *K*_0_ is a positive constant,

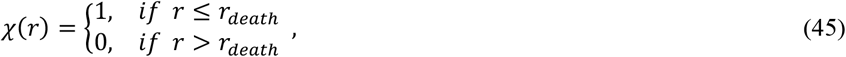

and then the expression for *P*_*death*_ simplifies to

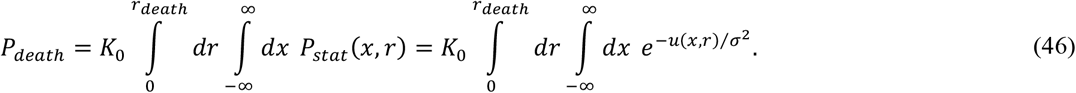

(Note the difference in the upper limit of integration over *r* from the previous formulas). Thus, we arrive at an intuitively understandable conclusion: the probability of a “Dickinsonia” dying per unit of time is proportional to the probability of its presence within the danger zone in the stationary distribution. This conclusion is analogous to the ergodicity condition (9), but unlike that condition, refers to the scenario of individual trajectories each interrupting at some time (different for different individual realizations of a stochastic trajectory).

We are aware that this derivation is not perfect and could be made stricter. In particular, the stationary distribution *P*_*stat*_ that appears in equations (43) and (46) was introduced as a function determined by *f*_1_ and *f*_2_ in the context of infinitely long stochastic trajectories [*x*(*t*), *r*(*t*)]. However, the death of organisms means that all the corresponding trajectories are finite. A more rigorous approach might involve a generalization of the Fokker-Planck equation, in line with equation (39), introducing additional new source and sink terms representing the birth and death of organisms. This would include terms accounting for certain, rather than probabilistic, predation within close distances. This direction seems overly complicated to us and might only hinders the analysis. As long as the probability *P*_*death*_ of being eaten is low, the stationary distribution *P*_*stat*_ should exist and be mainly determined by the dynamics of the organisms (primarily, the reaction of their nervous system and the work of effectors), with only a light tail in the region of *r* available to predators, justifying the derivation above.

### Appendix B. Calculation of the functional I for potential (23)-(26)

Based on equation (6), in the absence of neural activity, *f*_2_ can be expressed in terms of *u* and *A* as

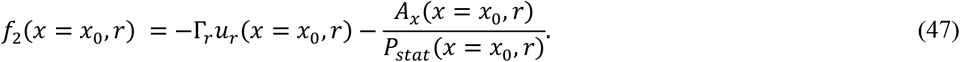

On the other hand, it equals the effective force arising from the nearest predator distribution, which was calculated in the previous paper^1^ [see also equation (3) here]:

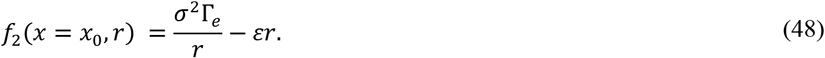

Taking into account equations (23)-(25) to compute *u*_*r*_, we arrive at the following condition:

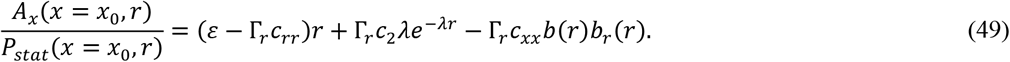

In general, the values of *A* at *x* ≠ *x*_0_ are not constrained, and various solutions would be compatible with condition (47) and the chosen form of the potential *u*. Here, we introduce an additional restriction that allows us to select one of these solutions: we assume that *Q* (and, therefore, *A*/*P*_*stat*_) does not depend on *x*. A possible motivation for this choice is as follows. We demonstrated in the previous paper that if the same value *λ* is used in the exponential functions in equations (25) and (26), and the prefactor before the logarithmic term in the potential *c*_*ln*_=1, and additionally if *C*_1_ = Γ_*r*_ λ_2_ *C*_2_ / *k*, then the function Δ(*r*) introduced in the previous paper^1^ vanishes, and the expression for *Q* simplifies to 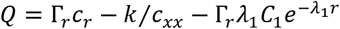, that is, does not depend on *x*.

With this assumption, all the dependence of *A* on *x* comes from the dependence of *P*_*stat*_ on *x*. In its turn, *P*_*stat*_ depends on *x* only via the *u*_1_ term, as equation (23) suggests, and therefore, *A* can be written as.

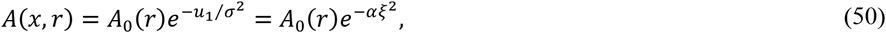

where we introduced a shorthand notation

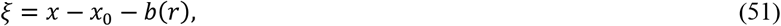

*α* = *c*_*xx*_/2*σ*^2^ as above, and *A*_0_(*r*) is some function of *r*. Then, the partial derivative

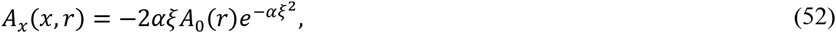

and its value at *x* = *x*_0_ is

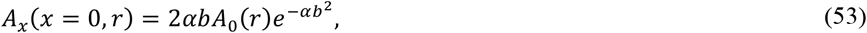

Then, by expressing *A*_0_(*r*) from equation (53), plugging it into equation (52), and taking into account equation (49), we get

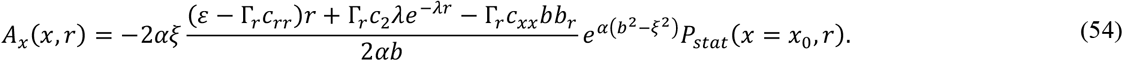

Taking into account that

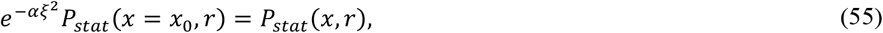

we arrive at an expression for *A*_*x*_/*P*_*stat*_ at arbitrary *x*.

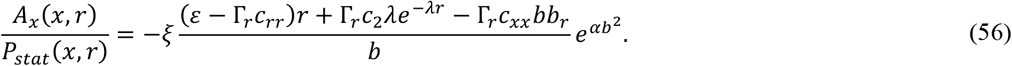

Then, the contribution to the velocity of the “Dickinsonia” due to the working nervous system can be computed as

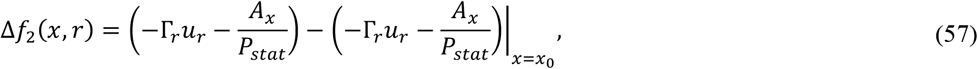

which, after boring mathematical derivations, simplifies to

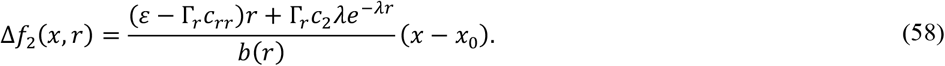

Therefore, a specific choice of the function *A* (based on the assumption that *Q* is independent on *x*) leads to a simple result valid for any numerical values of coefficients in the probe potential given by equation (23), namely, that the speed with which a “Dickinsonia” swims away from a predator is directly proportional to its nervous system excitation. Then, all integrals over *x* in the expression for the functional *I*, as seen in equation (18), can be determined analytically:

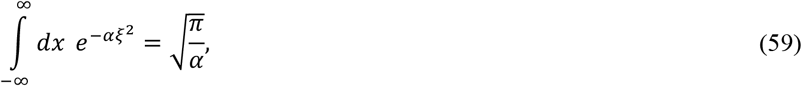

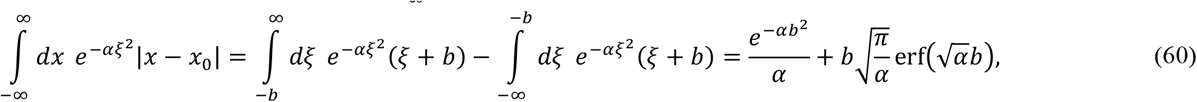

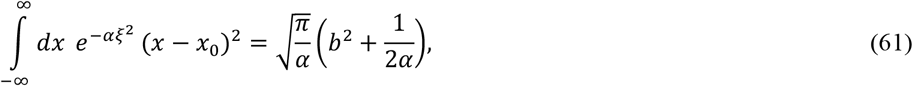

leading to equation (29) in the main text.

